# Mechanistic insights into B-cell activation and autoreactivity regulation in active SLE and remission

**DOI:** 10.64898/2026.02.27.708589

**Authors:** Yemil Atisha-Fregoso, Rita Pozovskiy, Meggan Mackay, Cynthia Aranow, Betty Diamond

**Author notes:** These authors equally contributed to this work (Yemil Atisha-Fregoso, Rita Pozovskiy).

## Abstract

**Objective:** To define cellular and molecular mechanisms distinguishing active systemic lupus erythematosus (SLE) from remission by profiling autoreactive antinuclear antigen– positive (ANA+) and non-autoreactive B cells subsets in three cohorts: active disease (SLE-A), long-term, drug free remission (SLE-R), and healthy controls (HC).

**Methods:** Peripheral blood B cells were phenotyped by flow cytometry, including ANA reactivity. Single-cell RNA sequencing (scRNA-seq) was performed on sorted ANA+ and ANA− subsets.

**Results:** Seven transcriptomic B cell clusters were resolved: quiescent (Naïve 1, Marginal Zone B cells [MZB], IgG Memory 1) and activated (Age-Associated B cells [ABCs], Naïve 2, IgM Memory, IgG Memory 2). SLE-A showed expansion of activated clusters, MZB contraction, and a higher IgG:IgM B cell ratio. SLE-R exhibited an “immunological reset,” distinct from healthy homeostasis, with reduced ABCs and IgG Memory 2, persistence of Naïve 2, and partial restoration of MZB and Naïve 1. Interferon-α (IFNa) signaling was elevated across clusters in SLE-A (SLE-A > SLE-R > HC), whereas TNF signaling was enriched in activated clusters across cohorts, with minimal differences between SLE-R and SLE-A. IFNa and TNF scores were inversely correlated. B cells predominantly expressed TNFR2, suggesting immunomodulatory TNF effects in remission. ANA+ cells in HC and SLE-R showed enriched FcγRIIb inhibitory and IL-4/STAT6 signaling, suggesting reinstated regulatory control.

**Discussion:** Compared to SLE-A, SLE-R was characterized by partial reversion to HC homeostasis with residual activation. These findings delineate immunologic features of remission and suggest therapeutic opportunities, including modulation of TNFR2, FcγRIIb, and IL-4 to help sustain remission.

**What is already known on this topic:** Some patients with SLE achieve complete clinical remission without treatment, referred as ‘immune reset’; the mechanisms that underlie this state have not been well characterized. Healthy individuals and patients with Systemic Lupus Erythematosus (SLE) normally harbor similar frequencies of autoreactive B cells; the checkpoints that regulate activation of these cells are not fully defined.

**What this study adds:** B cells, stratified by their reactivity to nuclear antigens (ANA), from active SLE (SLE-A), drug-free remission (SLE-R), and healthy controls (HC) were analyzed using single cell sequencing and flow cytometry. We identified B cells states associated with disease activity; SLE-R displayed a distinct profile that differed from SLE-A and HC. TNF signaling was present in activated B cell subsets in SLE-A and SLE-R. This persistence in SLE-R may reflect an immunomodulatory function of TNF on TNFR2, which is expressed on B cells. ANA+ cells in SLE-R and HC were enriched for inhibitory FcγRIIb and IL-4/STAT6 programs.

**How this study might affect research, practice or policy:** The signatures identified help define the “immunological reset” state in patients with SLE-R. We also identified pathways, such as type I IFN, TNFR2, FcγRIIb, IL-4/STAT6 as potential targets for maintaining remission.

## Introduction

B cells play a central role in Systemic lupus erythematosus (SLE) pathogenesis through autoantibody production, antigen presentation, and cytokine secretion. The success of B cell depletion by CD19-targeting CAR-T cells in selected patients with refractory SLE highlights the central role of B cells in disease pathogenesis [1]. IgG antinuclear antibodies (ANA) are part of the classification criteria for SLE and some are pathogenic [2]; thus, autoreactive plasma cells, particularly those producing ANA, are a hallmark of SLE. Our previous studies have shown that healthy individuals harbor ANA+ mature B cells at frequencies comparable to those observed in patients with SLE [3]. Despite this, healthy individuals do not produce pathogenic IgG ANA. This paradox raises critical questions about how ANA+ B cells are differentially regulated in health and disease and whether they are regulated differently from ANA-B cells. Recent work by our group and others has demonstrated the loss of regulatory checkpoints in SLE resulting in fewer anergic ANA+ B cells [4] and more IgG+ plasmablasts (PB) [5], suggesting abnormal activation of B cells, specifically of ANA+ B cells; however, the phenotypic and transcriptional differences between ANA+ and ANA-B cells across different B cell subsets have not been determined. Some patients with SLE achieve long-term remission, defined here as sustained absence of clinical activity without immunosuppressive medication for 3 years; use of hydroxychloroquine (HCQ) is permitted. This definition is consistent with, and more stringent than current definitions of remission, which might permit continued use of other immunosuppressants [6, 7]. This remission state, currently termed an “immunological reset”, is one in which autoantibodies and autoreactive B cells may persist without driving inflammation or tissue damage. The mechanisms of immune homeostasis that characterize this state have not been thoroughly studied. We studied a cohort of such patients (SLE in remission; SLE-R); and compared them to patients with active SLE (SLE-A) and healthy controls (HC) using flow cytometry and scRNA-seq to interrogate mechanisms of immune quiescence and regulation of pathogenic B cell responses.

## Materials and Methods

### Patients and healthy controls

Patients with SLE (American College of Rheumatology or Systemic Lupus International Collaborating Clinics criteria for SLE [8, 9]) were recruited from the Northwell Health Lupus Center of Excellence. Disease activity was assessed using the SLE Disease Activity Index-2K (SLEDAI-2K) [10]. Active SLE was defined as a SLEDAI-2K score ≥ 5, while SLE remission was defined as the absence of clinical activity for at least 3 years following a history of previous disease activity, with patients receiving no medication other than hydroxychloroquine (HCQ). All subjects provided written informed consent. The study was approved by the Northwell Health Institutional Review Board.

### Identification of autoreactive B cells and flow cytometry-based cell sorting

From each patient, 40cc of blood were obtained by venipuncture and peripheral blood mononuclear cells (PBMC) were isolated by gradient density centrifugation and cryopreserved at -80° C until analysis.

ANA+ B cells subsets were identified by flow cytometry using biotinylated nuclear and fluorochrome labeled antibodies and streptavidin, following the protocol previously described [4]. For cell sorting six B cell subpopulations were analyzed, stratified by ANA status (Supplementary Fig. 1A and 1B). To perform flow cytometry-based cell sorting, we identified 3 B cell subsets, further divided by ANA+ and ANA-: 1) CD27-CD38int/low B cells, containing mostly naïve B cells and CD27-IgD-cells which include ABCs; 2) CD27+ IgG+ memory B cells and 3) CD27+ IgM+ (unswitched) memory B cells. We also identified and quantified circulating PBs (not sorted). An example plot of ANA positivity is shown in supplementary Fig. 1C. Similar number of cells of each subpopulation from each participant were isolated to ensure sufficient representation of rare ANA+ subsets and increase the power to detect differences between ANA+ and ANA-B cells.

#### Single Cell (sc)RNA-seq analysis

Following quality control filtering, scRNA-seq profiles of 7587 B cells (3,167 from HC, 2464 from SLE-A and 1956 from SLE-R) were corrected for technical batch effects introduced by 10x Genomics sequencing runs using Harmony [11] followed by two-dimensional Uniform Manifold Approximation and Projection (UMAP) visualization.

To compare differential expression and pathway analyses in ANA+ and ANA-B cells in each cohort, we performed gene set enrichment analysis (GSEA) using MAST [12]. Using this framework, we performed a hurdle generalized linear model to identify differentially expressed genes (DEGs) between ANA+ and ANA-B cells within each cohort adjusted by cluster, to account for the different distribution of B cells into clusters. Genes that were identified as significantly differentially expressed in ANA+ B cells in each cohort were subjected to pathway enrichment analysis using Hallmark and Reactome gene sets.

A detailed description of statistical analyses and methods are provided in supplementary methods.

### Data availability

All sequencing data included in this study will be available upon publication at GEO NCBI database.

### Patient and Public Involvement statement

Patients or the public were not involved in the design or conduct of this research. The results of this study will be available to patients with SLE from the Feinstein Institutes upon request.

## Results

### Distinct B cell subset distribution and class switching patterns in SLE-A and SLE-R

We performed flow cytometric analysis (gating strategy is shown in Supplementary Fig. 1B) of PBMCs from 10 HC, 8 SLE-A, and 6 SLE-R (clinical characteristics are shown in Supplementary Table 1). We included in the analysis B cells associated with autoimmunity and aging in mice (ABCs). There is no standardized definition for these cells; we used CD19+, CD11c+, and CXCR5+, consistent with current literature [13]. Prior reports show majority of those cells are T-bet+ and ZEB2+, additional markers of ABCs [14, 15].

Naïve B cells were the predominant subset across cohorts, and significantly more abundant in SLE-R (Figs. 1A and 1B). Within CD27+IgM+ B cells, we observed two subsets: MZB (defined by CD1c and IgD) and IgM Memory (Supplementary Fig. 1B). Transitional B cells and PBs were significantly increased in SLE-A vs HC; IgG+ ABCs were increased in some SLE-A. MZB cells were reduced in SLE-A and SLE-R vs HC. The MZB:IgM Memory ratio was reduced overall in SLE and significantly lower in SLE-A vs HC (Fig. 1C). PCA of subset differences showed cohorts separation (Figs. 1D and 1E). PC1 was driven by transitional, DN1, ABCs (IgD+ and IgG+) and PBs; PC2 partially resolved HC vs SLE-R, driven by MZB, IgG Memory, and IgM Memory in HC, and Naïve B cells in SLE-R.

**Figure 1.**
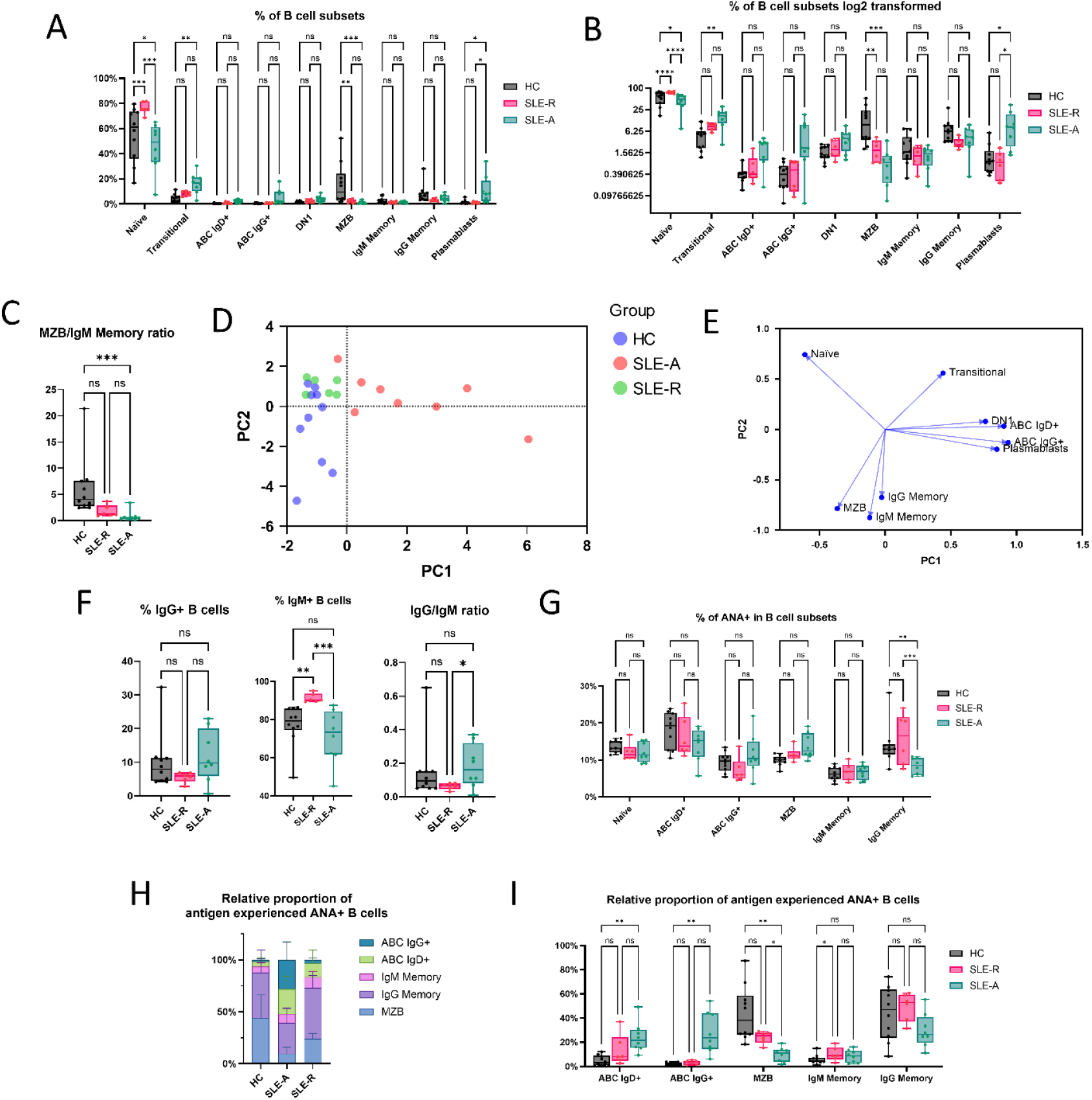
Subset distribution analysis by flow cytometry. (A) Boxplot shows the percentage of cells in each B cell subset across cohorts. (B) As in panel A but showing a log2-transformed y-axis to improve visualization of small values. (C) Boxplot comparing the MZB/IgM memory ratio between cohorts. (D) Scatterplot and (E) loading plot of the first two principal components of a principal component analysis of B cell subset frequency across cohorts. (F) Boxplots of the relative frequencies of IgG and IgM B cells within total B cells, and the IgG/IgM ratio for each cohort. (G) Boxplots of the relative frequency of ANA+ B cells within B cell subsets. (H) Stacked bar plot showing the relative B cells subset composition of antigen experienced ANA+ B cells across cohorts. (I) Boxplot showing the comparison of the relative proportion of each subset within the antigen experienced ANA+ B cell pool between different cohorts. Statistics. Comparisons were performed using the restricted maximum likelihood (REML) method in a mixed effects model and adjusted for multiple comparisons according to the Tukey-Kramer method (* p<0.05; ** p<0.01). MZB, Marginal Zone B; ANA, antinuclear antigen.

We previously demonstrated increased differentiation to IgG PBs [16] and serum IgG:IgM ratio in a subset of patients with SLE with more active disease [17]. We therefore quantified IgM+ and IgG+ B cells and their ratio (Fig. 1F). SLE-R had more IgM+ cells than SLE-A and HC; the IgG/IgM B-cell ratio was higher in SLE-A than in SLE-R. Additionally, serum from 11 HC, 8 SLE-A, and 6 SLE-R showed increased total IgG, ANA IgG, and anti-dsDNA IgG in SLE-A vs SLE-R and HC (Supplementary Fig. 1D).

### Differential distribution of ANA+ B cells in SLE-A and SLE-R

We next analyzed the relative percentage of ANA+ B cells in B cell subsets. For most subsets there was no difference in the percentage of ANA+ B cells across cohorts (Fig. 1G), which is consistent with our previous reports [4, 16]. We did, however, observe differences in the percentage of ANA+ cells between patient cohorts in IgG Memory B cells; SLE-A had the lowest percentage of ANA+ IgG Memory B cells.

We were interested in the relative distribution of the antigen-experienced ANA+ B cell pool, i.e. MZB, IgM Memory, IgG memory, and ABCs (IgD+ and IgG). In HC, ANA+ B cells were mostly concentrated in MZB and IgG Memory B cells (Fig 1H). In SLE-A, compared with HC, there was an increased proportion of ANA+ B cells among ABCs, both IgG+ and IgD+, and a reduction of the ANA+ B cells in MZB (Fig. 1H and I). Compared to SLE-A, patients with SLE-R have a higher percentage of ANA+ MZB. Interestingly, the proportion of ANA+ IgG ABCs is not different between HC and SLE-R, although the IgG+ ABC subset is small in both cohorts. There is, however, a higher percentage of ANA+ B cells in the IgM Memory subset of SLE-R compared to HC.

### Transcriptomic analysis identifies activated and quiescent clusters of B cells

We analyzed the transcriptomes of ANA+ and ANA-B cell subsets: 1) CD19+ CD27-CD38int (Including Naïve B cells and ABCs), 2) CD19+ CD27+ IgM+ (Including MZB and IgM Memory B cells) and 3) CD19+ CD27+ IgG+ (IgG Memory) from 8 HC, 6 SLE-R, and 8 SLE-A (sorting strategy in supplementary Fig. 2A). All HC and SLE-A participants were female; one SLE-R patient was male. SLEDAI-2K scores in the SLE-A cohort ranged from 5 to 14. Detailed demographics, clinical and laboratory data, and treatment information are summarized in Supplementary Table 2. The frequency of the sorted B cell subsets and ANA+ B cell prevalence are shown in Supplementary Fig. 2B.

Unsupervised clustering of the normalized, batch-corrected single cell transcriptomes identified seven distinct clusters (Fig. 2A). We assigned their identities in the UMAP embedding using canonical markers (Fig. 2B and 2C): Naïve B cells (2 clusters), lacking CD27 expression, and expressing IgM and IgD; unswitched memory B cells (USM; 2 clusters), defined by CD27 and IgM; and switched memory B cells (SM; 2 clusters) characterized by CD27 and IgG, with low IgD and IgM. We also identified age-associated B cells (ABCs) characterized by a high expression of ZEB2, ITGAX (CD11c) and TBX21 (T-bet) with low expression of CD27 and IgD. Although not directly sorted, ABCs were included within the CD27-population. IgD+ and IgG+ ABCs clustered together. As in flow cytometry, USM showed heterogeneity. One subset of MZB expressed high IgD and CD1c, consistent with circulating MZB [18]; the other subset lacked these markers, representing conventional IgM memory cells. Most clusters within each cohort included cells from multiple donors, except Naïve 1 and MZB in SLE-R (discussed later) and Naïve 1 in SLE-A, which derived from a single patient (Supplementary Fig. 2C). Cluster identities were consistent with the sorted populations (Supplementary Fig. 2D). The top 5 genes defining each cluster are represented in a heatmap (Fig. 2C). Naïve 1 and Naïve 2 were closely related as were IgM Memory and MZB and IgG Memory 1 and IgG Memory 2 B cells. ABCs clustered with both memory subsets (Fig. 2D), supporting a memory phenotype.

**Figure 2.**
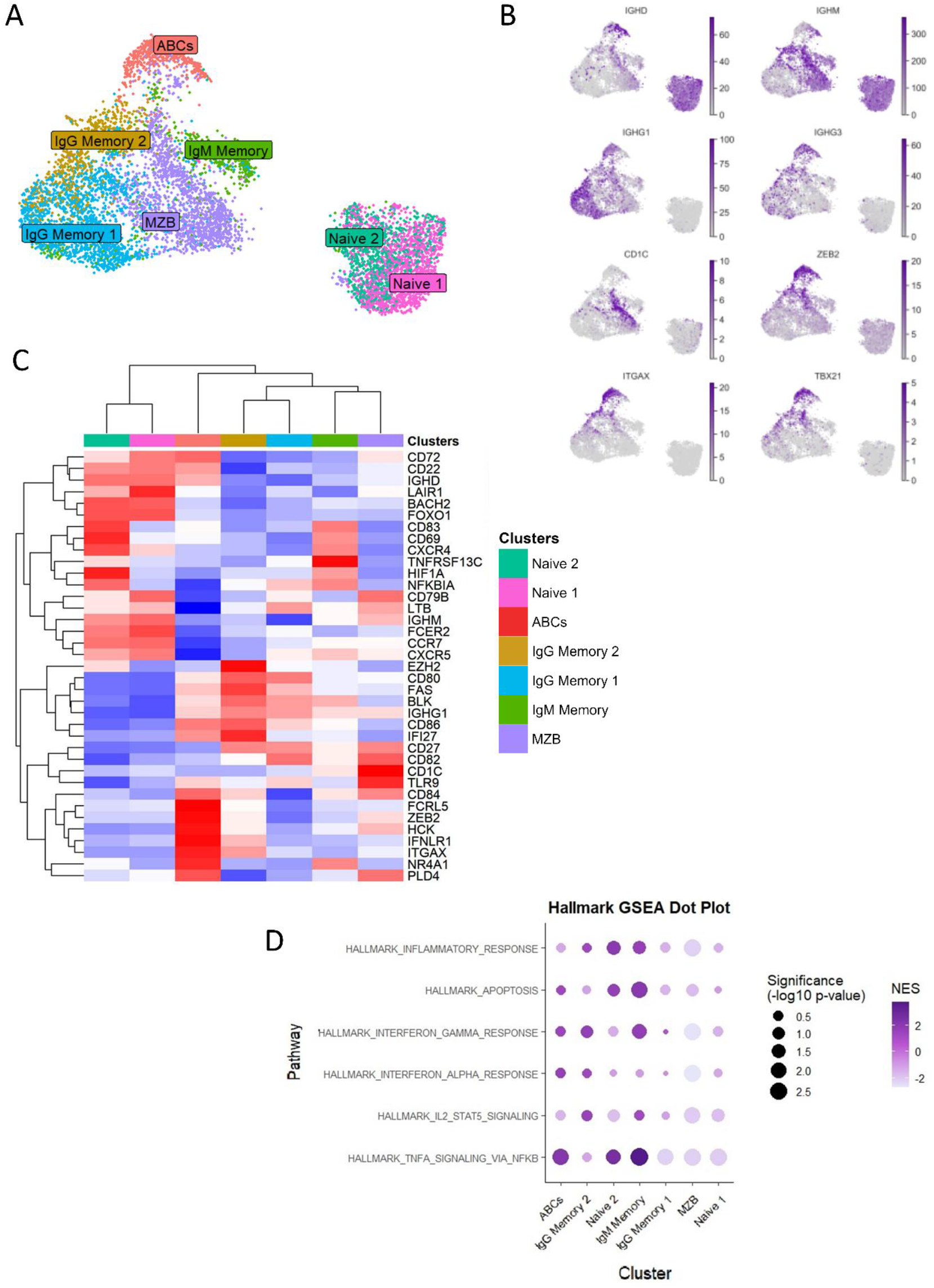
Cluster characterization in scRNA-seq. (A) UMAP visualization showing seven identified clusters. (B) UMAP feature plots display the expression of key markers used for B cell subset characterization. (C) Heatmap of relative expression for relevant B cell markers across clusters, with hierarchical clustering of both markers and clusters. (D) GSEA indicating pathway activation in selected clusters. UMAP, Uniform Manifold Approximation and Projection; GSEA, gene set enrichment analysis.

To explore the functional heterogeneity of B cell subsets, we performed Hallmark Gene Set Enrichment Analysis (GSEA) on the seven clusters. Overall, enrichment profiles showed activation signatures across ABCs, IgG Memory 2, Naïve 2 and IgM Memory subsets (Fig. 2E), although some heterogeneity was observed. Naïve 1, MZB and IgG Memory 1 clusters did not display upregulation of activation pathways, supporting a quiescent phenotype.

### Different distribution of B cells clusters between HC, SLE-R and SLE-A

B cell cluster distribution varied significantly across HC, SLE-A, and SLE-R (Fig. 3A). B cells from HC belonged exclusively to quiescent clusters: Naïve, MZB, and IgG Memory. SLE-A shifted toward activated clusters: ABCs, Naïve 2, IgM Memory, and IgG Memory 2.

**Figure 3.**
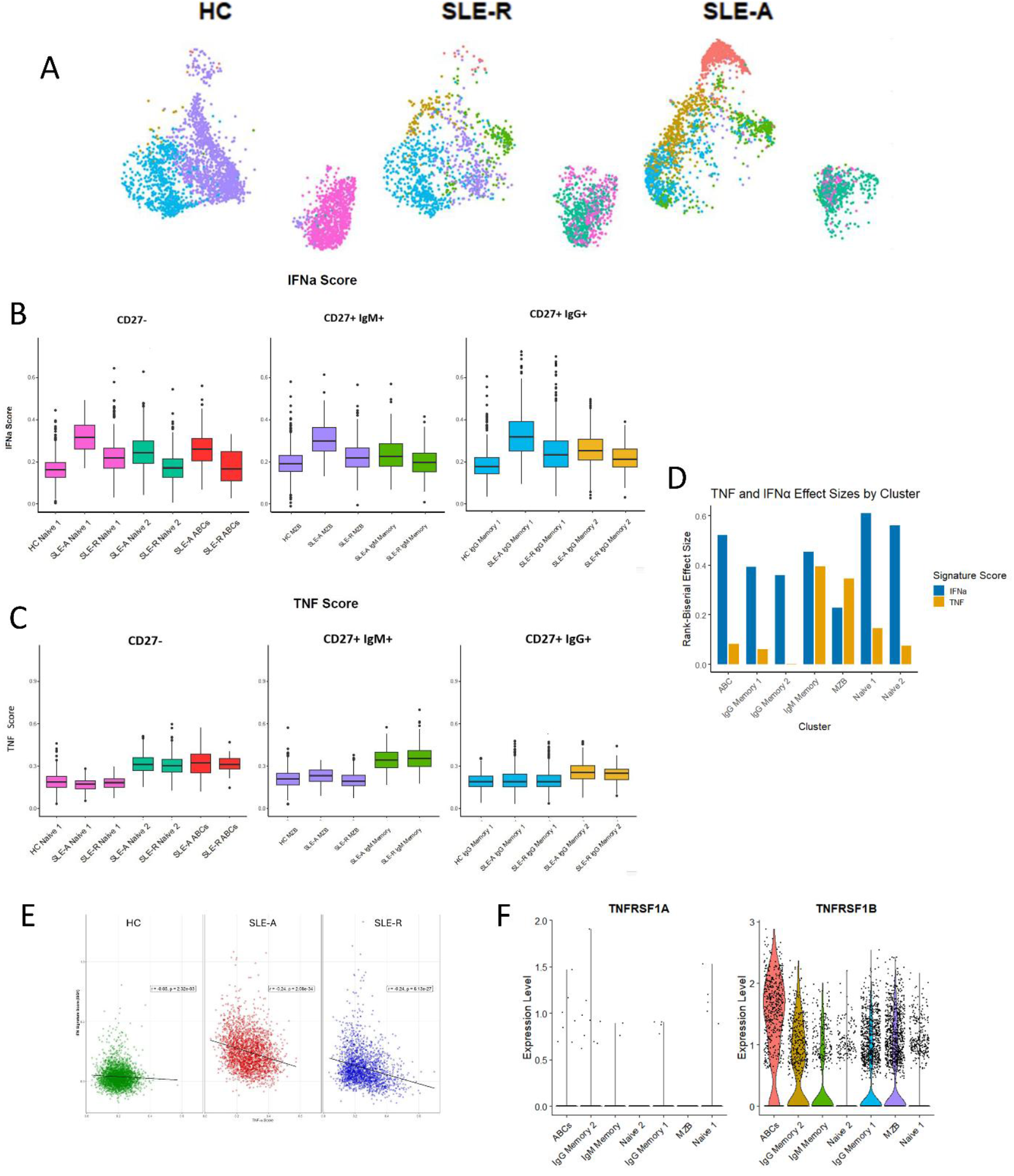
Cohort-specific signatures in B-cell scRNA-seq. (A) UMAP showing the distribution of cells by cohort of origin. (B) IFNa pathway scores across clusters stratified by cohort. (C) TNF pathway scores across clusters stratified by cohort. (D) Effect size for the comparisons between SLE-A and SLE-R for IFNa and TNF for each cluster (E) Correlation between IFNa and TNF pathway scores at the single-cell level within each cohort. (F) Expression of the principal TNF receptors (TNFRSF1A/TNFR1 and TNFRSF1B/TNFR2) across B-cell clusters from all subjects. UMAP, Uniform Manifold Approximation and Projection; IFNa, interferon alpha; TNF, tumor necrosis factor.

SLE-R had a unique profile. Compared to SLE-A, SLE-R showed lower proportions of ABCs and IgG Memory 2, a modest reduction in IgM Memory and higher proportions of Naïve and MZB cells. Interestingly, SLE-R displayed greater proportions of Naïve 2 B cells than SLE-A perhaps suggesting a tolerance checkpoint distinct from HC. In SLE-R, more MZB cells and Naïve B cells were observed in patients with serologically inactive (normal complement C3 and C4 and negative for anti-dsDNA antibodies) disease, while serologically active (low complement or anti-dsDNA antibodies) patients had a higher proportion of Memory 2 (Supplementary Fig. 3A).

### IFN and TNF are predominant activation pathways with different behavior in SLE-A and SLE-R

Pathway analysis showed enrichment of genes within IFN and TNF pathways in SLE-A and SLE-R. We examined IFNa downstream signaling using an IFNa score based on upregulated genes in B cells after in vivo exposure to IFNa [19] (Fig. 3B). IFNa scores were elevated across all clusters in SLE-A compared with SLE-R. In the non-activated clusters (present in all cohorts), we observed SLE-A > SLE-R > HC, and in activated clusters SLE-A > SLE-R, confirming heightened IFNa signaling as a hallmark of active SLE, while indicating residual IFNa activation in SLE-R.

We also observed TNF pathway gene enrichment in the GSEA and therefore, we derived a TNF activation score based on genes upregulated in B cells following TNF stimulation [19]. In contrast to IFNa, TNF scores were higher in activated clusters (Naïve 2, ABCs, IgM Memory and IgG Memory 2) and relatively similar between same subsets across cohorts (Fig. 3C). To quantify between-cohort differences for SLE-A and SLE-R, we calculated effect sizes for IFNa and TNF scores comparisons across clusters (Fig. 3D). IFNa effect sizes were higher than 0.3 for most clusters (except MZB, markedly reduced in SLE-A). Meanwhile, TNF showed minimal differences between SLE-A and SLE-R, with an effect size of more than 0.2 only in IgM memory and MZB (Fig. 3D). Interestingly, we observed an inverse correlation between TNF and IFNa signaling in SLE-A and SLE-R (Fig. 3E).

We observed that B cells predominantly express TNF receptor 2 (TNFR2; gene TNFRSF1B, also known as CD120b or p75), which modulates immune responses and tissue repair; with negligible expression of TNF receptor 1 (TNFR1; gene TNFRSF1A, also known as CD120a or p55), which is primarily responsible for mediating inflammatory responses and cell death (Fig. 3F).

### ANA+ B cells exhibit distinct subset distributions and pathway activation profiles across cohorts

To characterize autoreactive B cell frequency across clusters, we analyzed ANA+ and ANA− distributions across B cell subsets by cohort. Because cell distribution was determined by the sorting strategy, we compared predominant ANA+ and ANA− clusters within sorted groups (CD27−, CD27+ IgG, CD27+ IgM) (Fig. 4A). HC were excluded because each sorted group contained only one subset. In SLE-A, ABCs were significantly enriched within ANA+ cells in the CD27− compartment, with a significant reduction in Naïve B cells. SLE-A also showed a non-significant increase of ANA+ cells in IgG Memory 2. In SLE-R, IgG Memory 2 were significantly enriched within ANA+ cells. These findings indicate preferential accumulation of ANA+ cells in IgG Memory 2 and ABCs, and preferential differentiation of ANA+ cells into ABCs in SLE-A.

**Figure 4.**
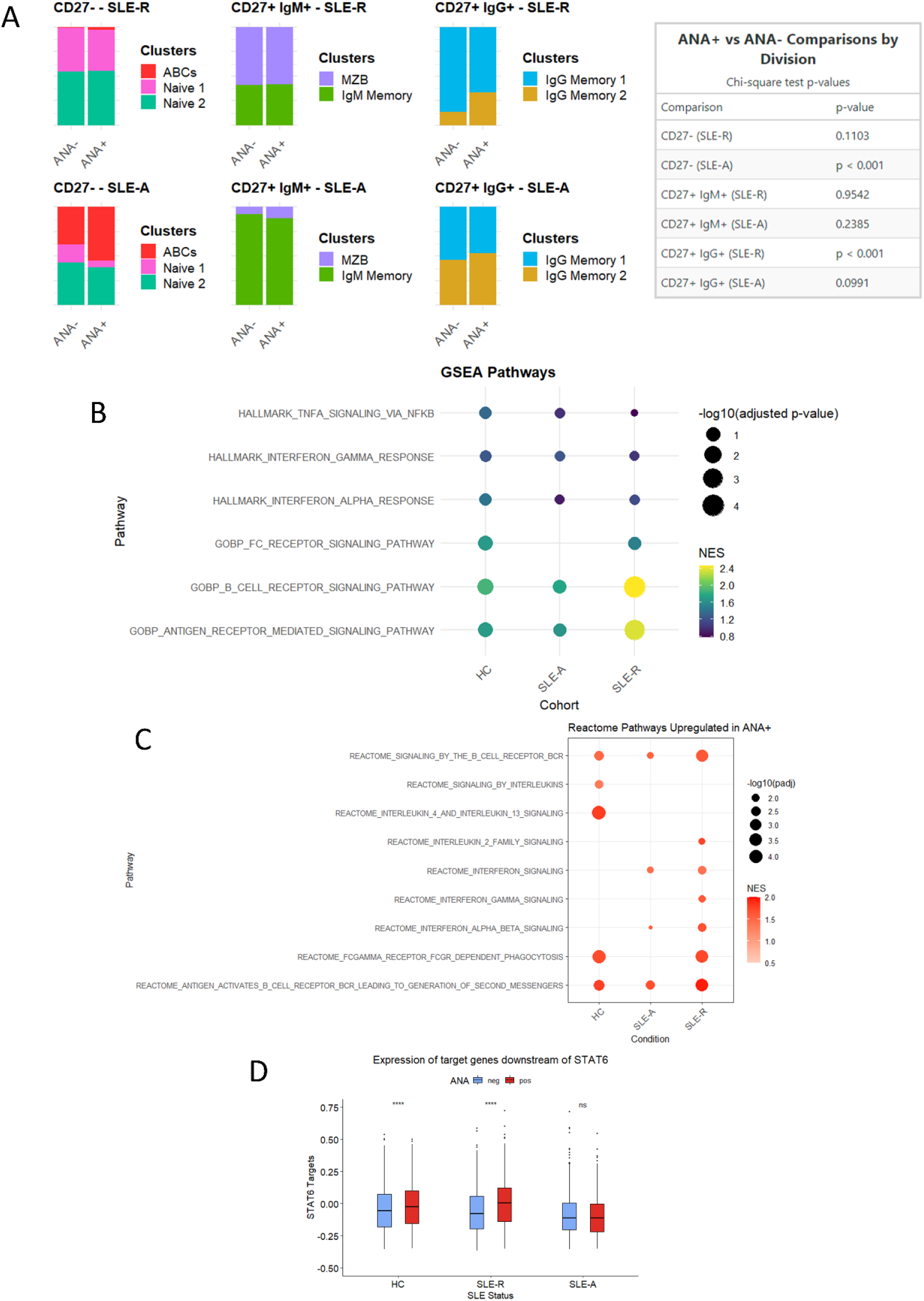
Differences between ANA+ and ANA– B cells across cohorts. (A) Stacked bar plots showing the relative distribution of B-cell subsets within CD27–, CD27+IgM+, and CD27+IgG+ B cells in SLE-A and SLE-R cohorts, stratified by ANA status (ANA+ vs ANA-). Fisher’s exact tests were performed within each sorting gate and cohort to compare subset proportions between ANA+ and ANA- groups. (B) GSEA of MSigDB Hallmark and GOBP pathways enriched in ANA+ versus ANA- B cells in each cohort. (C) GSEA of Reactome pathways upregulated in ANA+ versus ANA- B cells in each cohort. (D) Expression of STAT6 downstream target genes in ANA– and ANA+ B cells by cohort. Comparisons were made using the Wilcoxon rank-sum test (∗ p<0.05; ∗∗∗ p<0.001). ANA, antinuclear antigen; SLE-A, systemic lupus erythematosus active; SLE-R, systemic lupus erythematosus in remission; GSEA, gene set enrichment analysis; GOBP, Gene Ontology Biological Process; STAT6, signal transducer and activator of transcription 6.

We performed a GSEA of DEGs between ANA+ and ANA-B cells within each cohort (Fig. 4B and 4C). ANA+ B cells from all cohorts showed enrichment for signaling through the B cell receptor, which strongly suggests that these cells were exposed to antigen in vivo. Fc gamma receptor signaling pathway was enriched in SLE-R and in HC, but not in SLE-A. The only Fc gamma receptor expressed in B cells is FcγRIIb, an inhibitory receptor.

Notably, one of the strongest signals was IL-4/IL-13, enriched in ANA+ B cells in HC. As an immunomodulatory role of IL-4 in SLE has been suggested, we sought additional evidence of this pathway activation in ANA+ B cells. Because IL-4 signaling depends on STAT6, we examined expression of STAT6 downstream targets. SLE-A had lower expression of genes downstream STAT-6 than HC and SLE-R. Compared to their ANA-counterparts, ANA+ B cells from HC and SLE-R upregulated STAT6-dependent genes, suggesting IL-4 exposure (Fig. 4D). These findings pinpoint molecular pathways linked to sustained SLE activity or remission via specific modulation of ANA+ B cells.

## Discussion

This study provides a comprehensive characterization of B cell and autoreactive B cell heterogeneity in SLE across disease states, revealing distinct transcriptional programs and cellular phenotypes associated with disease activity and remission.

We identified seven distinct B cell clusters with varying activation states. Non-activated or quiescent clusters (Naïve, MZB, and IgG Memory) which predominate in HC, and activated clusters (ABCs, IgG Memory 2, IgM Memory, and Naïve 2) that characterize SLE-A. The ABC population comprises both IgM and IgG cells and expresses canonical ABC markers, including ZEB2, ITGAX (CD11c), and TBX21 (T-bet) [20]. The Naïve 2 subset shows evidence of activation; however, it is distinct from the “activated naïve B cells” described by Sanz and colleagues [21]. The previously described activated naïve B cells share features with ABCs (and DN2) and, in our dataset, would be encompassed within the ABC cluster rather than the Naïve 2 cluster, which does not express characteristic ABC markers such as T-bet, CD11c and ZEB2 and may not be poised to differentiate to ABCs.

SLE-R exhibits a unique B cell profile, indicating a different “homeostasis” compared to HC and demonstrating that “immune reset” may not signify reversion to the HC B cell profile. SLE-R demonstrated reduced proportions of ABCs, IgM memory and Memory 2 B cells compared to SLE-A, suggesting these subsets, together with PBs, may serve as cellular biomarkers of disease activity and may be implicated in pathogenesis. In flow cytometry, SLE-R showed a increased proportion of Naïve B cells. In line with our observations, many patients treated with CD19-targeting CAR-T cells display a high proportion of Naïve B cells and fewer memory B cells after reconstitution [22], although it is not known if they have a Naïve-2 like subpopulation. Maintaining a naïve B cell phenotype may help to prevent disease activity as it correlates with a lower IgG:IgM B cell ratio and a lower IgG:IgM ratio in serum. This has clear implications for disease activity because IgM ANA protect against the inflammatory effects of apoptotic debris, whereas IgG ANA are proinflammatory.

MZB are disproportionately reduced in SLE-A. It is possible that this reflects class-switch recombination in this population, consistent with the increased IgG:IgM B cell ratio. In SLE-R, the proportion of MZB cells is partially restored, as is the IgG:IgM B cell ratio. We also observed more class-switching to IgG in SLE-A compared to SLE-R. These findings align with our previous reports that an altered IgM to IgG ratio for anti-double stranded DNA antibodies is present in individuals at risk of developing SLE [23] and of B cell hyperactivation and hypergammaglobulinemia in SLE [17]. Additionally, they suggest restoration of MZB cells, which can differentiate into plasma cells producing protective IgM antibodies of [24], might be relevant for maintaining remission.

An intriguing observation is the persistence of Naïve 2 B cells in SLE-R, indicating that certain aspects of B cell dysregulation are present even during clinical remission and may be B cell-intrinsic in SLE, rather than driven by systemic inflammation. It is unclear whether Naïve 2 B cells contribute to the risk of flares or reflect an active homeostatic mechanism to maintain remission. Naïve 2 B cells were present in SLE-R with and without serological activity, while quiescent Naïve 1 B cells were observed only in patients without serological activity, suggesting activated Naïve 2 B cells may be on the road to disease activity rather than part of a homeostatic mechanism. It will be of interest to determine the activation state of Naïve B cells in individuals after B cell reconstitution following CAR-T cell therapy.

The IFN score is higher in SLE-R than in HC, although it is lower than in SLE-A. The persistence of an IFN signature was previously reported in patients with inactive SLE [25, 26], notably we observed it in patients without evidence of disease activity for years. A low IFN signature in blood and kidney cells of patients with lupus nephritis without a high activity index on biopsy but with high chronicity scores was recently reported [27], supporting that these changes are not limited to circulating cells and that the IFN signature is associated with immune cell activation and inflammation in tissues.

TNF scores were significantly elevated in activated B cell clusters, showing minimal differences between SLE-A and SLE-R within similar clusters. IFN and TNF signaling exhibited an inverse relationship, suggesting a complex interplay that may influence disease activity. A previous report described negative cross-regulation of TNF and IFN [28], mechanistically this was associated with plasmacytoid dendritic cells. Our findings in B cells suggest a potential regulatory loop of TNF and IFN in the context of SLE and B cell activation, or, alternatively, that cells were activated in different compartments with exposure to different cytokines.

The role of TNF in SLE is controversial. In SLE mouse models, deleterious and protective effect of TNF have been reported [29-32], with discrepancies possibly related to the stage of disease at the time of TNF exposure [32]. Increased TNF was previously reported in myeloid cells infiltrating the kidney in lupus nephritis [33]; however, low TNF predispose to SLE in both humans and mice [34]. Reports of small numbers of patients with SLE treated with TNF blockers for short periods of time, showed some improvement without flares [35]. Conversely, higher serum TNF has been reported in inactive disease when compared with active disease [36].

TNFR1 (TNFRSF1A) is the TNF receptor associated with a proinflammatory effect, while TNFR2 (TNFRSF1B) is associated with immunomodulatory functions in endothelial cells and regulatory T cells [34] and with B cells secreting IL-10 [37, 38]. Both receptors signal through NF-kB [39]; TNFR2 binds predominantly membrane-bound TNF whereas TNFR1 also binds soluble TNF, perhaps explaining why serum TNF levels do not correlate with B cell activation state. Predominant TNFR2 expression, together with sustained TNF pathway activation in B cells from SLE-R, suggests TNF may exert anti-inflammatory effects in B cell subsets in remission.

We observed a preferential accumulation of ANA+ B cells within IgG Memory 2 and ABC clusters in SLE-A, suggesting autoreactive B cells are more likely to undergo differentiation during active disease. This is consistent with evidence that ABCs represent a pathogenic population in autoimmunity and are enriched in autoreactivity [40, 41].

ANA+ B cells exhibit activation of proinflammatory pathways and BCR signaling, suggesting activation by autoantigens. Notably, we also observed enrichment of anti-inflammatory pathways, such as IL-4 and FcγRIIb, in ANA+ B cells from HC and SLE-R, but not SLE-A. Polymorphisms of FcγRIIb that reduce its function or expression are associated with SLE risk in humans [42], and B cell-specific deletion of FcγRIIb produces a lupus-like phenotype in mice [43].

The role of IL-4 in SLE is complex. Confirmation of STAT6 signaling in ANA+ B cells form SLE-R and HC provides evidence for a suppressive IL-4-mediated effect that may be diminished during active disease. While IL-4 can promote B cell proliferation and activation, facilitate class switching to IgG [44, 45] and reverse B cell anergy [46]; some reports show IL-4 can modulate autoimmunity [47] and inhibit the generation of ABCs and plasma cells, hallmarks of active SLE [48]. Our data suggest that activation of both FcγRIIb and IL-4 pathways likely contribute to regulate ANA+ B cell activation in HC and in SLE-R. These mechanisms appear compromised in SLE-A, and their restoration may reflect a tolerance mechanism in ANA+ B cells during remission.

Our previous data revealed a tolerance checkpoint controlling IgG plasma cell differentiation that is abrogated in SLE [5, 17]; the current data confirm this checkpoint, as SLE-R showed enrichment of ANA+ B cells in the activated IgG Memory 2 compartment without increase in PBs. We also identify a novel checkpoint present in SLE-R that blocks the differentiation of Naïve 2 B cells. High TNF signaling in this population may lead to elevated titters of IgM ANA which could be part of compensatory homeostasis. Future research should focus on longitudinal studies to better understand the mechanisms governing the transition to remission, the immunological changes preceding disease flares, and the development of therapies that can reinforce the regulatory pathways identified in this study.

Our findings demonstrate that remission in SLE does not simply represent a return to healthy immune homeostasis, but rather a distinct immunological state characterized by partial restoration of regulatory mechanisms and activation signatures such as TNF, FcγRIIb, and IL-4 that may be compensatory pathways in SLE-R, with the TNF pathway perhaps also modulating activation of B cells in SLE-A.

## Acknowledgements

We thank Arnon Arazi, Kaitlin Carroll, Alexandria Dalgish-Choi, Sophia Garcia, Houman Khalili, Cassandra Pond and Manami Watanabe for their valuable contributions to this work.

## Funding

This study was supported by the National Institutes of Health (NIH) grants 5P01AI172523 and 5U19AI144306, the Lupus Research Alliance (LRA) grant 2022GTSA-1211, and the Arthritis National Research Foundation (ANRF) grant 1068467.

## Notes

### Competing Interest Statement

The authors have declared no competing interest.

