## Supplementary Tables, Methods and Figures for "Mechanistic insights into B-cell activation and autoreactivity regulation in active SLE and remission"

Supplementary table 1. Clinical characteristics of subjects included in flow cytometry analysis.

| ID | Status | Race (ethnicity) | Gender | Age | Year of diagnosis | SLEDAI-2K | Clinical activity | DNA positive | Prednisone dose | HCQ dose | Immunosuppressants | Time inactive |
|---|---|---|---|---|---|---|---|---|---|---|---|---|
| F310 | SLE-A | W | F | 32 | 2014 | 14 | K / Se | 1 | 0 | 400 | MMF | NA |
| F695 | SLE-A | O | F | 43 | 2019 | 10 | MC / Ar / Fever | 0 | 0 | 400 | | NA |
| F729 | SLE-A | W(H) | F | 48 | 2004 | 10 | MC / Ar | 1 | 5 | 0 | | NA |
| F220 | SLE-A | B | F | 33 | 2013 | 14 | MC / Vasc | 1 | 6 | 200 | | NA |
| F135 | SLE-A | W(H) | F | 44 | 2010 | 8 | MC | 1 | 5 | 400 | MMF | NA |
| F502 | SLE-A | B | F | 40 | 2011 | 8 | Ar / AE | 1 | 10 | 400 | MMF | NA |
| F656 | SLE-A | B | F | 40 | 2015 | 5 | MC / Leu | 1 | 0 | 400 | | NA |
| F719 | SLE-A | B | F | 42 | 2019 | 8 | MC | 1 | 0 | 400 | MTX | NA |
| F659 | SLE-R | A | M | 27 | 2012 | 2 | NA | 1 | 0 | 400 | | 10 years |
| F134 | SLE-R | W | F | 53 | 2002 | 0 | NA | 0 | 0 | 400 | | 6 years |
| F474 | SLE-R | A | F | 38 | 2010 | 0 | NA | 0 | 0 | 400 | | 8 years |
| F504 | SLE-R | A | F | 36 | 2007 | 0 | NA | 0 | 0 | 400 | | 6 years |
| F094 | SLE-R | W(H) | F | 56 | 1999 | 0 | NA | 0 | 0 | 400 | | 10 years |
| F248 | SLE-R | W | F | 62 | 2011 | 0 | NA | 0 | 0 | 400 | | 5 years |
| F780 | HC | W | F | 36 | | | | | | | | |
| F788 | HC | W | F | 38 | | | | | | | | |
| F777 | HC | A | F | 62 | | | | | | | | |
| F759 | HC | W | F | 50 | | | | | | | | |
| F758 | HC | A | F | 30 | | | | | | | | |
| F523 | HC | A | F | 52 | | | | | | | | |
| F212 | HC | O | F | 50 | | | | | | | | |
| F760 | HC | W | F | 27 | | | | | | | | |
| F792 | HC | B | F | 38 | | | | | | | | |
| F776 | HC | W | F | 27 | | | | | | | | |

SLE-R=SLE in remission, SLA-A=SLE active, HC=Healthy control; Race(ethnicity): W=white, B=Black, A=Asian, Gu=Guyanese, O=Other, (H)=Hispanic; Gender: F=Female, M=Male; Clinical activity: MC=Mucocutaneous, K=Kidney, Se=Serositis, Leu=Leukopenia, AE=Angioedema, Ar=Arthritis, Vasc=Vasculitis; Immunosuppressants: MMF=Mycophenolate mofetil, MTX=Methotrexate. Time inactive: Time since recovery from last episode of activity if currently inactive.

Supplementary table 2. Clinical characteristics and demographic data of subjects included in scRNA-seq analysis.

| ID | Status | Race (ethnicity) | Gender | Age | Year of diagnosis | SLEDAI 2K | Clinical activity | DNA positive | Prednisone dose | HCQ dose | Immunosuppressants | Time inactive |
|---|---|---|---|---|---|---|---|---|---|---|---|---|
| F521 | SLE-R | W | F | 43 | 2006 | 4 | NA | 1 | 0 | 200 | | 8 |
| F445 | SLE-R | B | F | 37 | 2014 | 0 | NA | 0 | 0 | 400 | | >3 years |
| F442 | SLE-R | W (H) | F | 35 | 2009 | 0 | NA | 1 | 0 | 400 | | 9 years |
| F747 | SLE-R | W | F | 48 | 1990 | 0 | NA | 0 | 0 | 400 | | >10 years |
| F659 | SLE-R | A | M | 27 | 2012 | 0 | NA | 0 | 0 | 400 | | 10 years |
| F134 | SLE-R | W | F | 52 | 2002 | 0 | NA | 0 | 0 | 400 | | 4 years |
| B021 | SLE-A | B | F | 46 | 2002 | 6 | MC | 1 | 0 | 400 | | NA |
| F636 | SLE-A | B | F | 23 | 2019 | 6 | Se / Ly | 1 | 0 | 0 | | NA |
| F220 | SLE-A | B | F | 31 | 2013 | 14 | MC / Ar | 1 | 12.5 | 400 | MMF, Belimumab | NA |
| F761 | SLE-A | W (H) | F | 39 | 2001 | 10 | MC / K | 1 | 20 | 0 | MMF, Tacrolimus | NA |
| F656 | SLE-A | B | F | 39 | 2015 | 5 | MC | 0 | 0 | 400 | | NA |
| F155 | SLE-A | B | F | 39 | 2010 | 8 | Ar / Ly | 1 | 0 | 400 | Azathioprine | NA |
| F744 | SLE-A | W (H) | F | 51 | 1986 | 10 | MC / Ar / K | 1 | 5 | 400 | Voclosporin | NA |
| F721 | SLE-A | B | F | 40 | 1996 | 12 | MC / Vasc | 1 | 20 | 0 | MMF, Tacrolimus | NA |
| F212 | HC | O | F | 48 | | | | | | | | |
| F758 | HC | A | F | 27 | | | | | | | | |
| F759 | HC | A | F | 48 | | | | | | | | |
| F760 | HC | W | F | 25 | | | | | | | | |
| F757 | HC | A | F | 35 | | | | | | | | |
| F780 | HC | W | F | 34 | | | | | | | | |
| F788 | HC | W | F | 36 | | | | | | | | |
| F786 | HC | A | F | 30 | | | | | | | | |

SLE-R=SLE in remission, SLA-A=SLE active, HC=Healthy control; Race(ethnicity): W=white, B=Black, A=Asian, O=Other, (H)=Hispanic; Gender: F=Female, M=Male; Clinical activity: MC=Mucocutaneous, K=Kidney, Se=Serositis, Ly=Lymphadenopathy, Ar=Arthritis, Vasc=Vasculitis; Immunosuppressants: MMF=Mycophenolate mofetil. Time inactive: Time since recovery from last episode of activity if currently inactive.

### SUPPLEMENTARY FIGURE LEGENDS

**Supplementary Figure 1. Experiment design and ELISA results**. (A) Schematic representation of the study design. A different set of patients were included to perform scRNA-seq from sorted B cell subsets and flow cytometry on PBMCs. (B) Gating strategy used for flow cytometry analysis. (C) Representative scatterplot panel from flow cytometry data showing ANA+ on B cells. (D) Results from IgG ELISA (left), ANA IgG ELISA (middle) and dsDNA IgG ELISA performed in 6 SLE-R and 7 of 8 SLE-A included in scRNA-seq analysis and 11 independent HC.

scRNA-seq, single-cell RNA sequencing; PBMC, Peripheral blood mononuclear cells; ANA, antinuclear antigens; SLE-R, patients with systemic lupus erythematosus in remission; SLE-A, patients with systemic lupus erythematosus with active disease; HC, healthy controls.

**Supplementary Figure 2. Differences in sorted B cell subsets between groups by flow cytometry.** (A) Gatting strategy for flow cytometry-based cell sorting for scRNA-seq analysis. For each B cell subset ANA+ and ANA- B cells were sorted. (B) Distribution of B cell subsets and proportion of ANA+ cells within subsets for the samples included in scRNA-seq analysis. (C) Stack plots of the differential distribution of each B cell cluster for each individual patients grouped by cohort. (D) UMAP visualization shows the distribution of cells according to the subset of origin in cell sorting.

scRNA-seq, single-cell RNA sequencing; ANA, antinuclear antigens; UMAP, Uniform Manifold Approximation and Projection.

**Supplementary Figure 3. Cluster distribution in SLE-R according to SLEDAI** (A) UMAP visualization of distribution of cells coming from patients with SLEDAI-2K of 0 (serologically inactive) or SLEDAI-2K 2-4 (serologically active) in the SLE-R remission cohort.

UMAP, Uniform Manifold Approximation and Projection; SLE-R, patients with systemic lupus erythematosus in remission.

**Supplementary methods**

### Sample Collection and Cryopreservation

Peripheral blood samples from SLE patients and healthy controls (HCs) were collected in BD Vacutainer® Sodium Heparin Blood Collection Tubes (158 USP units, BD Biosciences, Cat# 367874; 9 mL × 4 tubes per donor). PBMCs were isolated within 4 hours of collection using SepMate™-15 tubes (StemCell Technologies) and Ficoll-Paque™ PLUS density gradient centrifugation. Whole blood was diluted 1:1 with HBSS and layered over Ficoll in SepMate tubes. After centrifugation at 1200 × g for 10 minutes, the PBMC layer was harvested, washed twice in HBSS, and resuspended in CryoStor® CS10 cryopreservation medium (BioLife Solutions, Cat# 210102). Cells were frozen at −80 °C in Mr. Frosty™ containers and transferred to liquid nitrogen for long-term storage.

### Identification of autoreactive B cells or Incubation with Biotinylated HeLa Nuclear Extract

ANA+ B cells were identified using biotinylated nuclear extract staining, following the protocol previously described [4]. In brief, cells were incubated with biotinylated nuclear extract (Bio-NE) prepared from nuclei of HeLa cells, diluted in HBSS containing 5% non-fat dry milk, for 30 minutes on ice. Following incubation, cells were washed twice with HBSS supplemented with 5% FBS and resuspended in a surface staining cocktail with streptavidin-APC included as part of the regular antibody panel to detect Bio-NE binding. For cell sorting for scRNA-seq, the cocktail included CD19 PE-Cy7 (BioLegend, 302216), CD27 PE (Invitrogen/eBioscience, 12-0279-42), CD38 PE-eFluor™ 610 (Invitrogen/eBioscience, 61-0389-42), IgG FITC (BD Biosciences, 555786), IgA PerCP-Cy5.5 (BioLegend, [Insert]), Streptavidin APC (BioLegend, 405207), CD1c BV421 (BioLegend, 331525), and Fixable Viability Dye eFluor™ 506 (Invitrogen/eBioscience, 65-0866-14). Cells were incubated with the antibody cocktail for 25 minutes on ice, protected from light. After staining, cells were washed with 2 mL of HBSS with 5% FBS, centrifuged at 1500 rpm for 5 minutes, and the supernatant was aspirated. Each sample was resuspended in HBSS buffer with 5% FBS and kept on ice. Collection tubes were pre-filled with X-VIVO™ 15 serum-free medium (Lonza, Walkersville/Whittaker BioProducts, 02-060Q) and kept at 4ºC until sorting. One-third of each sample was transferred to a separate tube for ANA⁻ sorting and the remaining two thirds were used for ANA+ sorting. All samples were maintained on ice until processing on a BD FACSAria™ cell sorter.

**Sequencing and Genomic Alignment**

Following FACS sorting, cells were fixed and permeabilized using the Next GEM Single Cell Fixed RNA Sample Preparation Kit (PN-1000414). Single-cell RNA sequencing libraries were then prepared using the Chromium Fixed RNA Kit, Human Transcriptome (PN-1000476), according to the manufacturer's protocol. The resulting FASTQ files were uploaded to the 10X Genomics cloud analysis platform. Within this platform, raw sequencing reads were aligned to the human genome (GRCh38-2020-A) using the Cell Ranger pipeline (10X Genomics), which generated gene expression matrices in HDF5 format for downstream analysis.

**Quality Control, Data Processing, Unsupervised analysis and Cluster Identification**

Using the Seurat (v5) package, filtered gene-by-cell matrices for 132 samples were merged and then processed using a standard unsupervised workflow (normalization, scaling, dimensionality reduction, batch correction, cell clustering, and differential gene expression analysis). First, the merged dataset was normalized and log-transformed (scaling factor = 10,000). The top 2,000 highly variable genes were identified using the variance stabilizing transformation method and used for scaling the data. During scaling, the percentage of mitochondrial gene expression was regressed out to mitigate unwanted variation. Dimensionality reduction was then performed with principal component analysis (PCA) over the top 2,000 variable genes, and the top 40 principal components (PCs) were selected for downstream analysis based on visual inspection of the elbow plot.

Following quality control filtering, scRNA-seq profiles of 7,587 B cells (3,167 from HC, 2,464 from SLE-A and 1,956 from SLE-R) were corrected for technical batch effects introduced by

10x Genomics sequencing runs using Harmony [1] followed by two-dimensional Uniform Manifold Approximation and Projection (UMAP) visualization. Cells were clustered using the Louvain algorithm implemented in the FindClusters Seurat function with a resolution parameter of 0.4.

To focus the analysis on B cell populations, infiltrating cells were removed from the dataset. Feature plots and violin plots were generated to visualize the expression of canonical marker genes for various cell types. Based on these expression patterns, clusters representing infiltrating cells such as T cells, monocytes, and NK cells were identified and removed by subsetting the Seurat object; remaining cells were reanalyzed to reconstruct stable clusters for visualization and subsequent analysis.

Remaining B cell clusters were annotated manually by visual examination of the expression of canonical markers, *CD27*, *CD38*, *IGHD*, *IGHM*, *IGHG1*, *IGHG3*, *CD1C*, *ITGAX*, and *TBX21*. Differential gene expression analysis was performed using the FindMarkers function in Seurat to identify genes enriched in each cluster.

**Heatmap construction**

To visualize the top representative genes that defined each cluster, we generated a heatmap using the dittoSeq R package. For this, we first calculated average expression values for each cluster using the AverageExpression function in Seurat, focusing on a set of manually selected marker genes chosen based on the FindMarkers output. These average expression values were then used to construct the heatmap, which displays the expression of these representative genes across all identified clusters.

**GSEA analysis**

To identify enriched biological pathways within each cell cluster, Gene Set Enrichment Analysis (GSEA) [2] was performed using the differentially expressed genes for each cluster that were identified using the FindMarkers function in Seurat. The resulting gene list for each cluster was then ranked based on the average log2 fold change (avg_log2FC) values. These ranked gene lists were used as input for GSEA, which was performed using the *fgsea* R package and the Hallmark gene set collection from MSigDB. GSEA results were then filtered to retain only pathways related to B cell function, lupus, SLE, TLR signaling, interferon signaling, TNF signaling, and interleukin signaling. Finally, to visualize the filtered GSEA results, a dot plot was generated using *ggplot2*, where dot size represents the adjusted p-value and dot color represents the Normalized Enrichment Score (NES).

**Per-Patient Cluster Distribution Analysis**

To examine the distribution of cell clusters across individual patients and disease cohorts, a bar plot was generated. Cell counts were calculated for each patient within each disease cohort and cell cluster. These counts were then normalized to represent the proportion of each cell cluster within each patient. Finally, a stacked bar plot was created using *ggplot2* to visualize these proportions, with each bar representing a patient, the bar height representing the total cell proportion (100%), and the colored segments within each bar representing the proportion of each cell cluster. The plot was divided by disease cohort to allow for comparisons between different patient groups.

**IFN and TNF scoring and boxplots**

To investigate the influence of interferon-alpha (IFNa) and tumor necrosis factor-alpha (TNF) signaling on B cell subsets across different disease cohorts, an immune response gene expression analysis (IREA) dictionary, published by the Hacohen lab [3], was used to compile a list of orthologous human genes upregulated in B cells after stimulation with IFNa and TNF. Module scores for IFNa and TNFa were calculated using Seurat's AddModuleScore function. For visualization purposes, cells were grouped based on cluster and disease cohort, and *ggplot2* was used to generate boxplots displaying the IFNa or TNFa module score for each combination of disease cohort (HC, SLE-A, SLE-R) and B cell subset. Wilcoxon rank-sum tests were conducted to compare IFNa and TNFa module scores (1) *within* each defined B cell subset across the different cohorts and (2) *between* B cell subsets within each cohort. To control for multiple comparisons, p-values from the Wilcoxon tests were adjusted using the Benjamini-Hochberg method. Effect sizes for each comparison were quantified using Cliff's delta, and these effect sizes (specifically, the rank-biserial correlation coefficient) were visualized in a bar graph using *ggplot2*. The bar graph displayed the effect size for each IFNa and TNFa signature score within each B cell subset, allowing for a direct comparison of the magnitude of effect between different conditions.

**Correlation Analysis of IFNa and TNFa Scores by B Cell Subset and Disease Status**

To explore the relationship between IFNa and TNF signaling within cohorts a scatterplot was generated using *ggplot2*, with each point representing a single cell. A linear regression smooth was added to each facet. To quantify the strength and statistical significance of the correlation within each facet, Pearson correlation coefficients and p-values were calculated. These values were then displayed as annotations on the scatterplot.

**Stackplot Visualization of Cluster Composition by ANA Status**

To visualize the relationship between B cell cluster composition and anti-nuclear antibody (ANA) status within each disease cohort, pairs of stacked bar plots were generated. First, cells were grouped by cohort and ANA status (positive or negative). To simplify the visualization, B cell clusters were further grouped into three categories: "CD27-" (including ABCs, Naïve 1, and Naïve 2 clusters), "CD27+ IgM+" (including MZB and IgM Memory clusters), and "CD27+ IgG+ " (including IgG Memory 1 and IgG Memory 2 clusters). Stacked bar plots were then created using *ggplot2*, with each bar representing a specific combination of SLE status and cluster group, and the segments within each bar representing the proportion of cells with positive or negative ANA status. The individual plots were combined into a single figure using the *patchwork* R package. To statistically assess the relationships within the SLE-A and SLE-R cohorts, chi-square tests were conducted. For each B cell subset division, the test compared the distribution of ANA-positive and ANA-negative cells between the individual clusters within that division in each cohort. Note that the HC cohort was excluded from the chi-square analysis due to the limited number of B cell clusters present within each division. The resulting p-values from these chi-square tests were then formatted into a publication-style table using the *gt* R package.

### MAST analysis

To compare differential expression and pathway analyses in ANA+ and ANA- B cells in each cohort, we performed gene set enrichment analysis (GSEA) using MAST [4]. Using this framework, we performed a hurdle generalized linear model to identify DEGs between ANA+ and ANA- B cells within each cohort adjusted by cluster (~ANA + Cluster. Genes that were identified as significantly differentially expressed in ANA+ B cells in each cohort were subjected to pathway enrichment analysis using Hallmark and Reactome gene sets. GSEA results were then filtered to retain only pathways related to B cell function, lupus, SLE, TLR signaling, interferon signaling, TNF signaling, and interleukin signaling. Finally, to visualize the filtered GSEA results, dot plots were generated using *ggplot2*, where dot size represents the adjusted p-value and dot color represents the Normalized Enrichment Score (NES).

### STAT6 Target Gene Expression Boxplot

To investigate upregulation of genes downstream of STAT6 a list of target genes known to be expressed in B cells (CCL17, CCL22, FCER2, IL4I1, ALOX15, EGR2, IRF4, XBP1) was curated based on evidence from two prior publications [5, 6]. Next, a STAT6 target gene expression signature score was calculated using the *AddModuleScore* function from Seurat. Finally, cells were grouped by cohort and ANA status, and a boxplot was created displaying the distribution of the score. Wilcoxon rank-sum tests were performed to compare the groups.

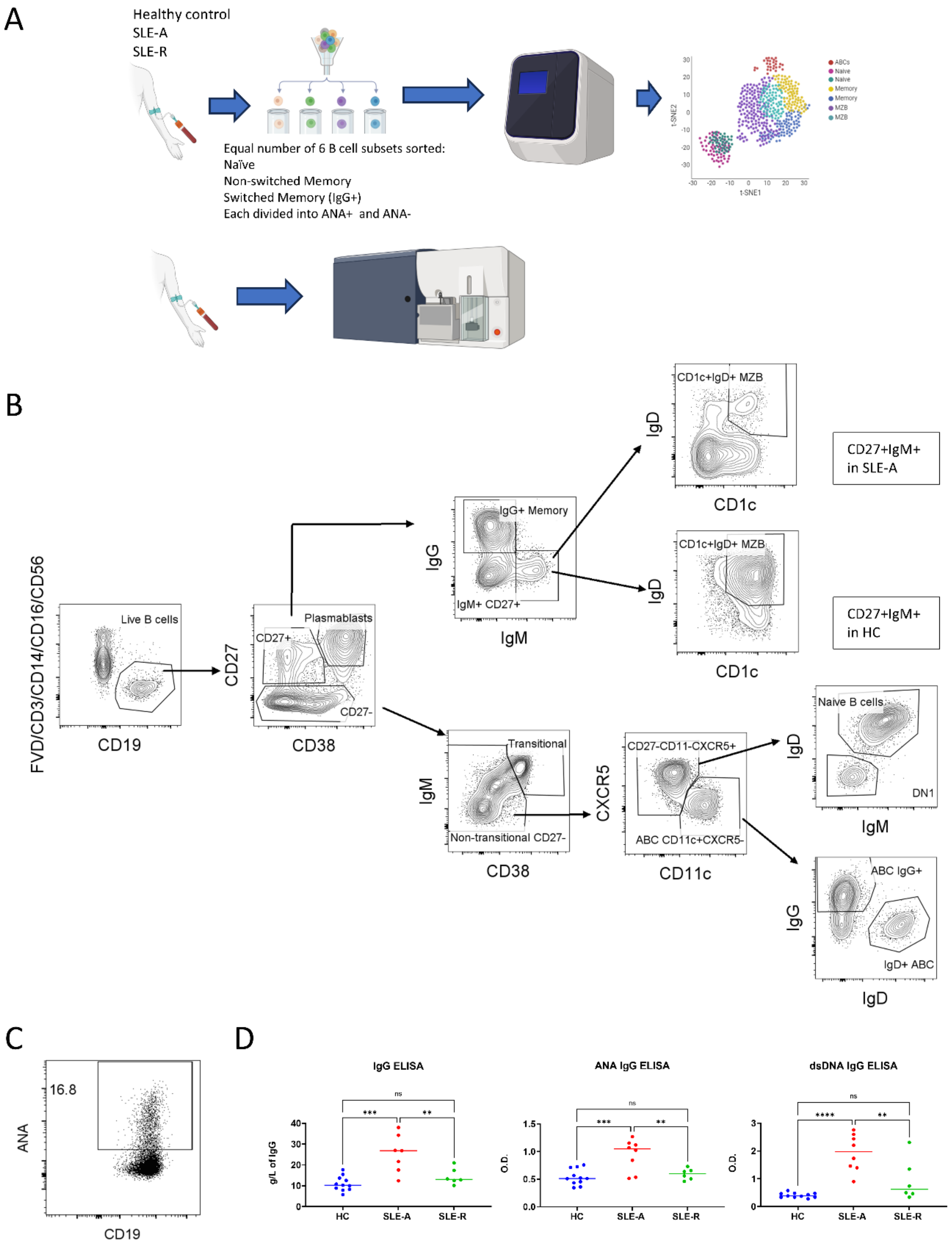
A
Healthy control
SLE-A
SLE-R
Equal number of 6 B cell subsets sorted:
Naïve
Non-switched Memory
Switched Memory (IgG+)
Each divided into ANA+ and ANA-
ABCs
Naive
Naive
Memory
Memory
MZB
MZB
t-SNE1
t-SNE2
B
FVD/CD3/CD14/CD16/CD56
Live B cells
CD19
CD27
Plasmablasts
CD27+
CD27-
CD38
IgG
IgG+ Memory
IgM+ CD27+
IgM
IgD
CD1c+IgD+ MZB
CD1c
CD27+IgM+ in SLE-A
CD1c+IgD+ MZB
CD1c
CD27+IgM+ in HC
IgM
Transitional
Non-transitional CD27-
CD38
CXCR5
CD27-CD11-CXCR5+
ABC CD11c+CXCR5-
CD11c
IgD
Naive B cells
DN1
IgM
IgG
ABC IgG+
IgD+ ABC
IgD
C
ANA
16.8
CD19
D
IgG ELISA
g/L of IgG
HC
SLE-A
SLE-R
ANA IgG ELISA
O.D.
HC
SLE-A
SLE-R
dsDNA IgG ELISA
O.D.
HC
SLE-A
SLE-R
ns
***
**
****

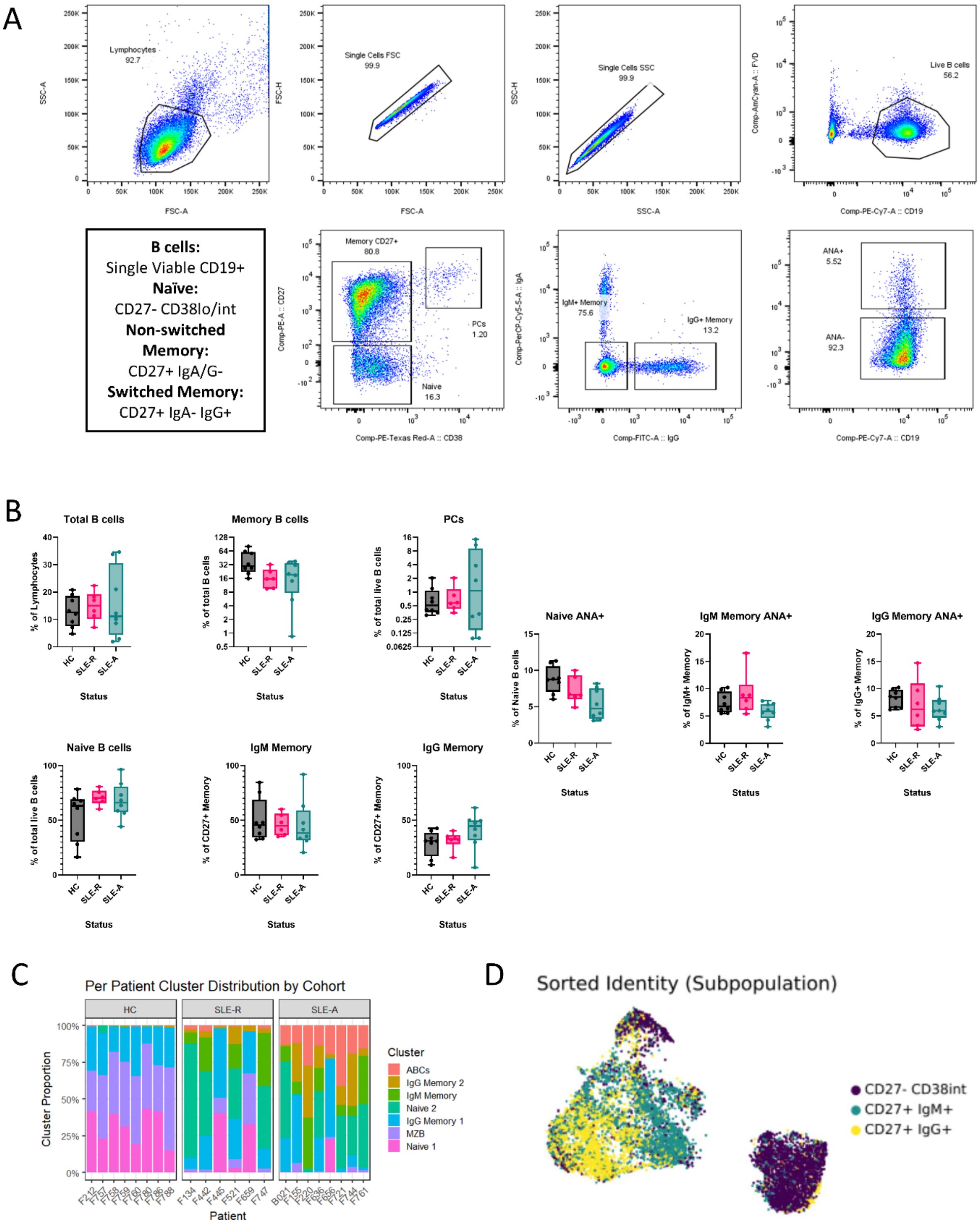

A
Lymphocytes
92.7
Single Cells FSC
99.9
Single Cells SSC
99.9
Live B cells
56.2
SSC-A
FSC-H
SSC-H
Comp-AmCyan-A :: FVD
FSC-A
SSC-A
Comp-PE-Cy7-A :: CD19
B cells:
Single Viable CD19+
Naïve:
CD27- CD38lo/int
Non-switched
Memory:
CD27+ IgA/G-
Switched Memory:
CD27+ IgA- IgG+
Memory CD27+
80.8
PCs
1.20
Naive
16.3
Comp-PE-A :: CD27
Comp-PE-Texas Red-A :: CD38
IgM+ Memory
75.6
IgG+ Memory
13.2
Comp-PerCP-Cy5-5-A :: IgA
Comp-FITC-A :: IgG
ANA+
5.52
ANA-
92.3
Comp-PE-Cy7-A :: CD19
B
Total B cells
% of Lymphocytes
Memory B cells
% of total B cells
PCs
% of total live B cells
Naive ANA+
% of Naive B cells
IgM Memory ANA+
% of IgM+ Memory
IgG Memory ANA+
% of IgG+ Memory
Naive B cells
% of total live B cells
IgM Memory
% of CD27+ Memory
IgG Memory
% of CD27+ Memory
HC
SLE-R
SLE-A
Status
C
Per Patient Cluster Distribution by Cohort
HC
SLE-R
SLE-A
Cluster Proportion
Cluster
ABCs
IgG Memory 2
IgM Memory
Naive 2
IgG Memory 1
MZB
Naive 1
Patient
D
Sorted Identity (Subpopulation)
CD27- CD38int
CD27+ IgM+
CD27+ IgG+

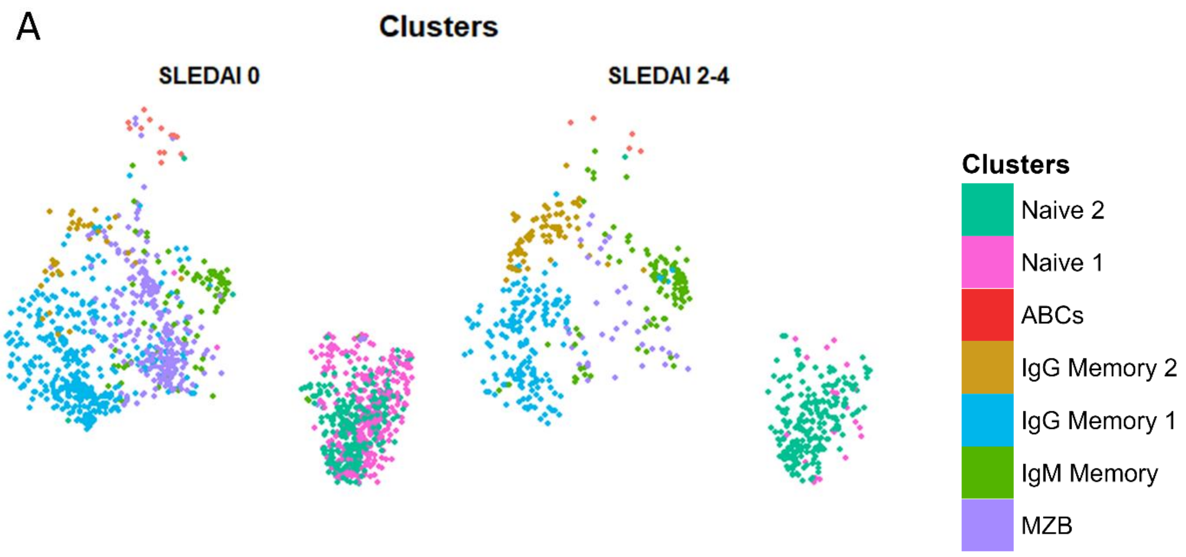

A
Clusters
SLEDAI 0
SLEDAI 2-4
Clusters
Naive 2
Naive 1
ABCs
IgG Memory 2
IgG Memory 1
IgM Memory
MZB
